# Denitrifying metabolism of the methylotrophic marine bacterium *Methylophaga nitratireducenticrescens* strain JAM1

**DOI:** 10.1101/180950

**Authors:** Florian Mauffrey, Alexandra Cucaita, Philippe Constant, Richard Villemur

## Abstract

*Methylophaga nitratireducenticrescens* strain JAM1 is a methylotrophic, marine bacterium that was isolated from a denitrification reactor treating a closed-circuit seawater aquarium. It can sustain growth under anoxic conditions by reducing nitrate (NO_3_^−^) to nitrite (NO_2_^−^), which accumulates in the medium. These physiological traits are attributed to gene clusters that encode two dissimilatory nitrate reductases (NarGHJI). M. *nitratireducenticrescens* strain JAM1 also contains gene clusters encoding two putative nitric oxide reductase (NO) reductases and one putative nitrous oxide (N_2_O) reductase, suggesting that NO and N_2_O can be reduced by strain JAM1. In this study, we show that strain JAM1 can reduce NO to N_2_O and N_2_O to N2 and can sustain growth under anoxic conditions by reducing N_2_O as the sole electron acceptor. Although strain JAM1 lacks a gene encoding a dissimilatory copper-(NirK) or cytochrome cd1-type (NirS) NO_2_^−^ reductase, NO_3_^−^-amended strain JAM1 cultures produce N_2_O, representing up to 6% of the N-input. NO_2_^−^ was shown to be the key intermediate of this production process. In NO_3_^−^amended cultures, we analyzed denitrification genes in succession of net N_2_O-production and -consumption phases at the gene expression level. These phases were found to correlate with changes in the expression levels of the NO reductase gene *cnorB1* and *nnrS*, which indicated NO production in the cultures.

**Importance:** By showing that all the three denitrification reductases are active, this demonstrates that *Methylophaga nitratireducenticrescens* JAM1 is one of many bacteria species that maintain genes associated primarily with denitrification, but not necessarily related to the maintenance of the entire pathway. The reason to maintain such incomplete pathway could be related to the specific role of strain JAM1 in the denitrifying biofilm of a denitrification reactor from which it originates. The small production of N_2_O via NO in strain JAM1 did not involve Nar contrary to what was demonstrated in *Escherichia coli. M. nitratireducenticrescens* JAM1 is the only reported *Methylophaga* species that has the capacity to grow under anoxic conditions by using NO_3_^−^ and N_2_O as sole electron acceptors for its growth. It is also one of a few marine methylotrophs that is studied at the physiological and genetic levels in relation to its capacity to perform denitrifying activities.

## Introduction

The complete denitrification pathway describes the successive reduction of nitrate (NO_3_^−^) to nitrite (NO_2_^−^), nitric oxide (NO), nitrous oxide (N_2_O), and nitrogen (N2) (1). This process is used by bacteria for respiration in environments with low oxygen concentrations and with NO_3_^−^ as an electron acceptor. The process is driven by metalloenzymes NO_3_^−^ reductase, NO_2_^−^ reductase, NO reductase, and N_2_O reductase (2). As a facultative trait, denitrification occurs frequently across environments and is performed by bacteria of diverse origins (3). However, numerous bacterial strains have been isolated with incomplete denitrification pathway, meaning that at least one reductase-encoding gene cluster is missing. As proposed by Zumft (3), the four steps of reduction from NO_3_^−^ to N2 could be seen as a modular assemblage of four partly independent respiratory processes that respond to combinations of different external and internal signals. This could explain the vast diversity of bacteria with incomplete denitrification pathway that can sustain growth with one of the four nitrogen oxides as electron acceptor. Another purpose of the incomplete pathway is related to detoxification, as nitrite and NO are deleterious molecules (4-7).

*Methylophaga nitratireducenticrescens* JAM1 is a marine methylotrophic gammaproteobacterium that was isolated from a naturally occurring multispecies biofilm that has developed in a methanol-fed, fluidized denitrification system that treated recirculating water of the marine aquarium in the Montreal Biodome (8, 9). This biofilm is composed of at least 15 bacterial species and of numerous protozoans (10, 11), among which *Methylophaga* spp. and *Hyphomicrobium* spp. compose more than 70% of the biofilm (12). Along with the denitrifying bacterium *Hyphomicrobium nitrativorans* NL23, *M. nitratireducenticrescens* JAM1 was shown to be the representative of the *Methylophaga* population in the biofilm (8).

*M. nitratireducenticrescens* JAM1 is considered as a nitrate respirer as it can grow under anoxic conditions through the reduction of NO_3_^−^ to NO_2_^−^, which accumulates in the culture medium (8). This trait is correlated with the presence of two gene clusters encoding dissimilatory nitrate reductases (*narGHJI,* referred as Nar1 and Nar2) in the genome of *M. nitratireducenticrescens* JAM1, which we showed that both contribute to NO_3_^−^ reduction during strain JAM1 growth (13). Anaerobic growth by strain JAM1 is a unique among *Methylophaga* spp. that were described as strictly aerobic bacteria (14). Genome annotation revealed that stain JAM1 seems to maintain an incomplete denitrification pathway with the presence of gene clusters encoding two putative cytochrome bc-type complex NO reductase (cNor) (*cnor1: norQDBCRE* and *cnor2: norCBQD)* and one putative dissimilatory N_2_O reductase (N_2_OR) (*nosRZDFYYL*), but lacks gene encoding a dissimilatory copper-(NirK) or cytochrome cd1-type (NirS) NO_2_^−^ reductase. These gene clusters have been shown to be transcribed (13). These data suggest that *M. nitratireducenticrescens* JAM1 has other respiratory capacities by performing NO and N_2_O reduction.

In this study, we aimed to assess the denitrification capacities of *M. nitratireducenticrescens* JAM1. Our results show that strain JAM1 can reduce NO to N_2_O and then to N2. It can use N_2_O as a source of energy for its growth under anoxic conditions. Through our investigation, we found that strain JAM1 cultured with NO_3_^−^ under anoxic and oxic conditions generates a small amount of N_2_O, despite the absence of gene encoding NirK or NirS. NO_2_^−^ was found to be a key intermediate of this production process. By using the JAM1 Δ*narG1-narG2* double mutant, we showed that the two Nar were not involved in N_2_O production via NO. We analyzed at the gene expression level the succession of N_2_O production and consumption with the denitrification genes *cnorB (cnorB1* and *cnorB2)* and *nosZ,* and also *nnrS* that encodes a NO-sensitive regulator. We found that gene expression level of *cnorB1* and *nnrS* increased during the N_2_O production phase, which suggest the presence of NO.

## Results

### M. nitratireducenticrescens JAM1 grows on N2O under anoxic conditions

Strain JAM1 was cultured under anoxic conditions with either NO_3_^−^ in the medium or with N_2_O injected in the headspace as the sole electron acceptor. Both types of culture received the same electron equivalent of NO_3_^−^ or N_2_O (1.3 mmole vial^-1^ or 18.2 and 36.4 mg-N vial^-1^, respectively) according to:

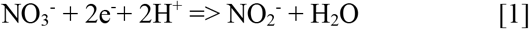

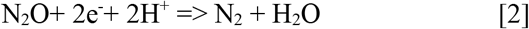

In N_2_O-amended cultures, N_2_O decrease was apparent from the start and consumption continued for 48 hours (Fig. 1A). The N_2_O decrease paralleled strain JAM1 growth with almost complete N_2_O consumption. The NO_3_^−^-amended cultures showed complete NO_3_^−^ consumption and equivalent nitrite accumulation after 24 h (Fig. 1B). However, slower growth than that recorded for the N_2_O cultures was observed. Such growth kinetics could be related to the toxicity of nitrite that accumulated in the medium. Both types of culture reached equivalent biomass concentration (t test on the last 4-time points, P >0.05).

**Figure 1:**
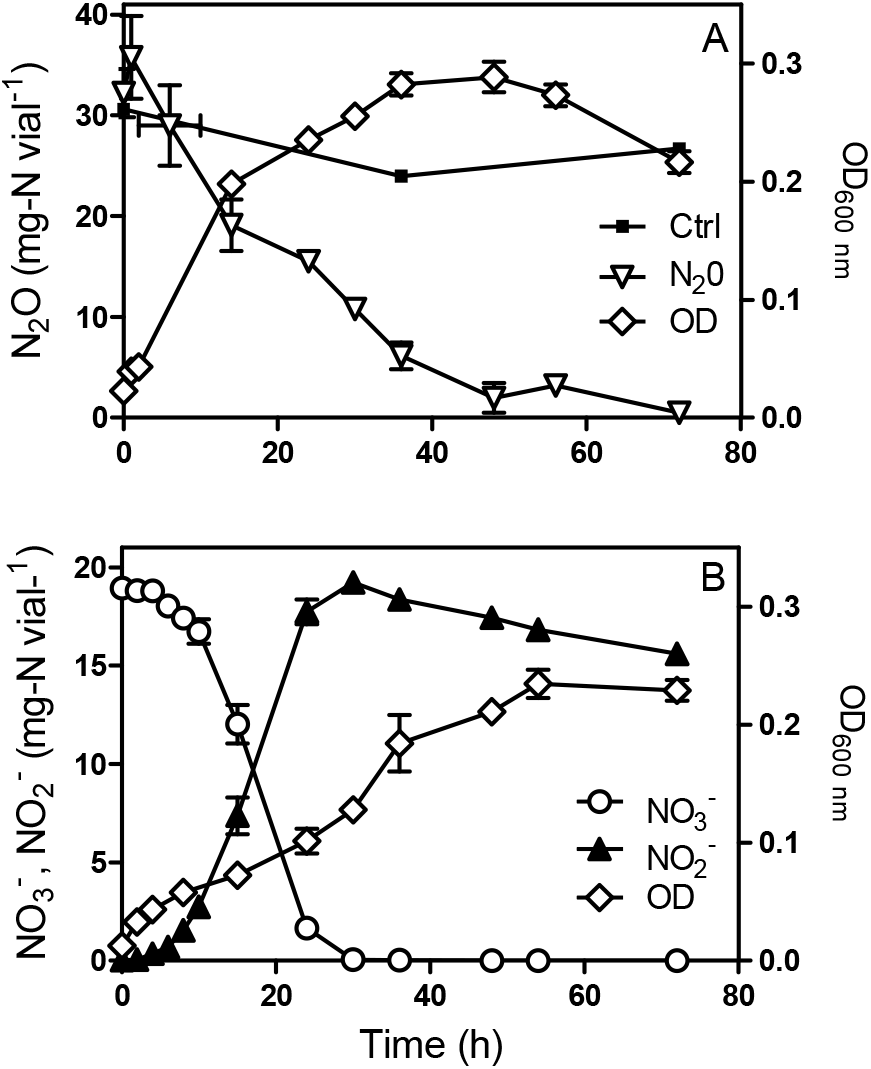
Methylophaga nitratireducenticrescens JAM1 growth with N_2_O or NO_3_^−^ as an electron acceptor. Strain JAM1 was cultured with 36.4 mg-N vial^-1^ N_2_O (A) or 18.2 mg-N vial^-1^ NO_3_^−^ (B) under anoxic conditions. N_2_O, NO_3_^−^ and NO_2_^−^ concentrations and growth were measured over different time intervals. Control (A): N_2_O injected in noninoculated vials. To minimize oxygen contamination, sampling was performed using a glove bag inflated with nitrogen gas. Data represent mean values ± standard deviation (SD; n=3).

### M. nitratireducenticrescens JAM1 consumes N_2_O under oxic conditions

In a previous study, we demonstrated that strain JAM1 can consume NO_3_^−^ under oxic growth conditions with equivalent accumulation of NO_2_^−^ (13). Culturing strain JAM1 under oxic conditions with N_2_O (3.5 mg-N vial^-1^) also showed a complete N_2_O consumption within 24 h (Fig. 2). In presence of O2, the strain JAM1 oxic cultures reached higher (4-5-times) biomass concentration than the anoxic cultures. N_2_O-amended cultures yielded equivalent biomass (O.D. ~1.2) than NO_3_^−^-amended oxic cultures (20 mg-N vial^-1^) (Fig. 2).

**Figure 2.**
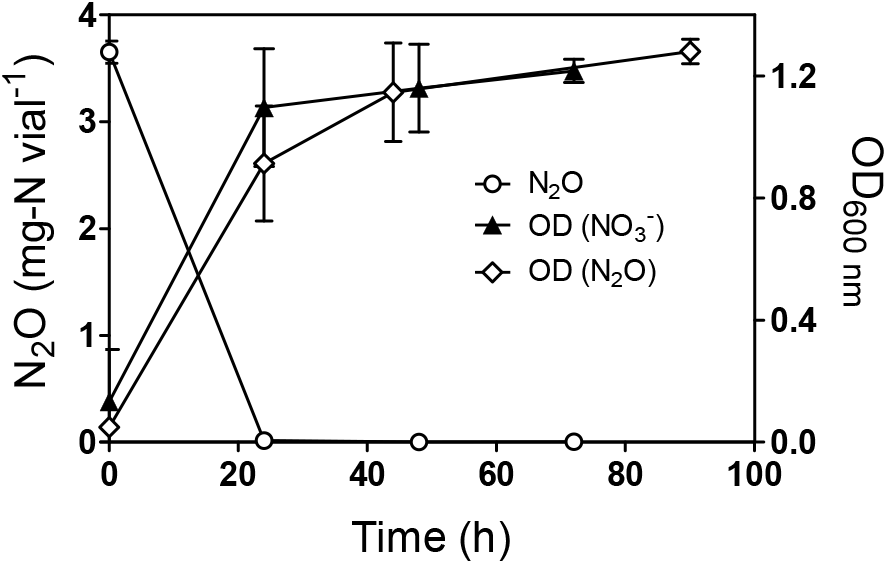
N_2_O consumption by *Methylophaga nitratireducenticrescens* JAM1 under oxic conditions. Strain JAM1 was cultured with 3.5 mg-N vial^-1^ N_2_O or 22 mg-N vial^-1^ NO_3_^−^ under oxic conditions. N_2_O and growth were measured over different time intervals. Data represent mean values ± SD (n=3).

### N_2_O production in NO_3_^−^-amended cultures

During the first assays to test the capacity of strain JAM1 to reduce N_2_O, cultures were performed with N_2_O (3.5 mg-N vial^-1^) but with the addition of NO_3_^−^ (20 mg-N vial^-1^) to make sure that growth would occur. Although N_2_O was completely consumed within 24 h, a net production of N_2_O was observed after 48 h (data not shown). The production of N_2_O by strain JAM1 is puzzling, as its genome does not contain NirS or NirK.

To further investigate this observation, strain JAM1 was cultured under anoxic conditions with NO_3_^−^, and NO_3_^−^, NO_3_^−^ and N_2_O were measured (Fig. 3A). Complete NO_3_^−^ reduction (19.3 ± 0.3 mg-N vial^-1^) was performed within 55 h. The nitrite level reached 17.5 ± 0.2 mg-N vial^-1^ over this period and decreased slowly to 15.9 ± 0.5 mg-N vial^-1^. N_2_O production initiated when NO_3_^−^ was nearly reduced and reached 0.70 ± 0.21 mg-N vial^-1^ after 55 h of incubation (Fig. 3A). N_2_O was completely reduced after 127 h. In parallel, for cultures in which the headspace was flushed with argon, N2 production was also measured. The corresponding results show an increase of N2 in the headspace (Fig. 3A) by 1.14 ± 0.54 mg-N vial^-1^ after 127 h, which represent 6.0 ± 2.9% of the N input.

Under oxic conditions, NO_3_^−^ reduction (17.4 ± 2.1 mg-N vial^-1^) was complete after 24 h with equivalent NO_2_^−^ accumulation (17.1 ± 1.3 mg-N vial^-1^). N_2_O accumulation started after complete nitrate reduction (Fig. 3B) and increased to reach 0.31 ± 0.32 mg-N vial^-1^ after 96 h of incubation (1.7% of N input). Unlike trends observed for the anoxic cultures, no N_2_O consumption was observed in the oxic cultures.

**Figure 3.**
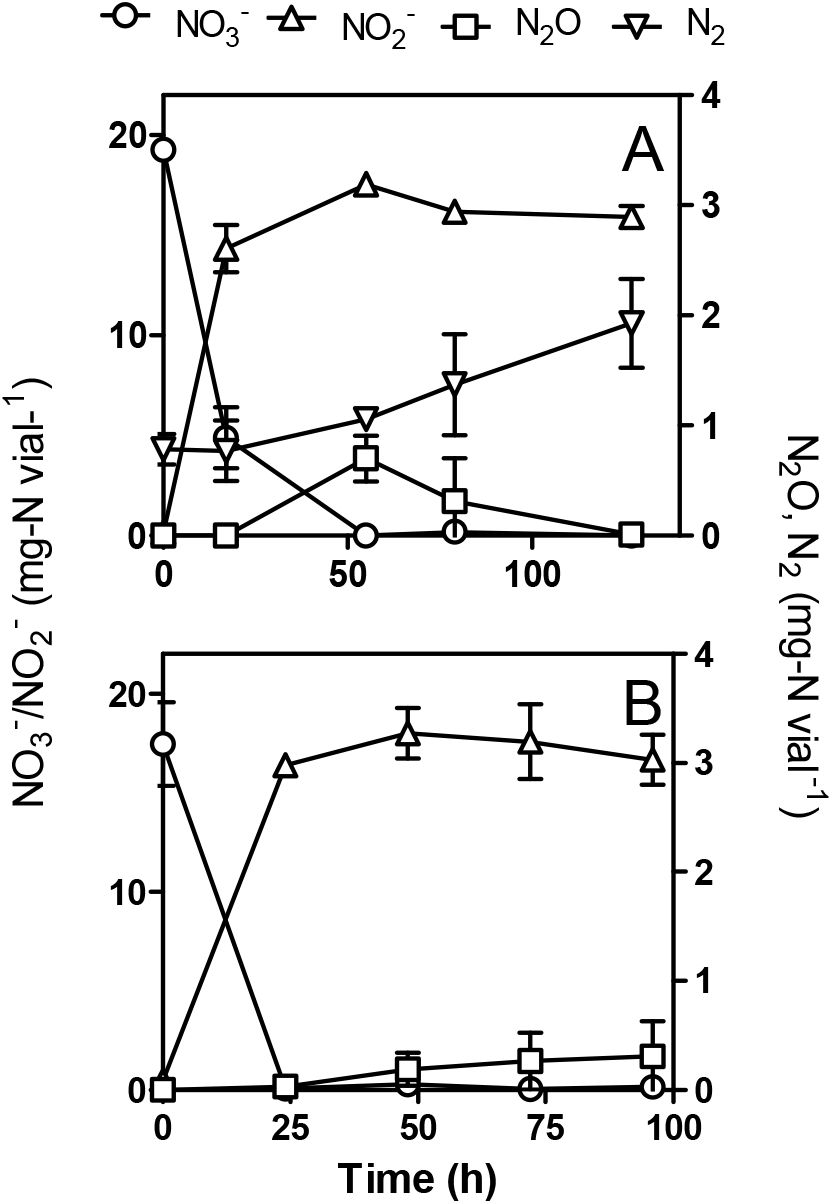
N_2_O production by *Methylophaga nitratiredu-centicrescens* JAM1. Strain JAM1 was cultured under anoxic (A) or oxic (B) conditions with NO_3_^−^ (22 mg-N vial^-1^). NO_3_^−^, NO_2_^−^, N_2_O and N2 (panel A only) concentrations were measured over different time intervals. Data represent mean values ± SD (n=3).

### Influence of ammonium on NO_3_^−^ and NO_2_^−^ consumption, and N_2_O production and consumption by M. nitratireducenticrescens JAM1

The original 1403 medium recommended by the ATCC for culturing *Methylophaga* spp. contains 20.9 mg-N vial^-1^ NH4Cl and 0.1 mg-N vial^-1^ ferric ammonium citrate (see Material and Methods). Therefore, NO_3_^−^ transformation in NH_4_^+^ should not be necessary in *Methylophaga* metabolism for nitrogen assimilation in biomass (Fig. S1). For the next set of experiments, we aimed to determine the effect of the absence of NH_4_^+^ on the dynamics of NO_3_^−^ and NO_2_^−^ consumption, and N_2_O production and consumption. We hypothesized that that forcing strain JAM1 to reroute some NO_3_^−^ for N assimilation would affect denitrification and thus growth rates. Strain JAM1 was cultured with *ca.* 20 mg-N vial^-1^ NO_3_^−^ under anoxic or oxic conditions in NH4Cl-free medium (Fig. 4A and B). Growth pattern observed under anoxic conditions was similar between the regular and NH4Cl-free cultures, as also the growth pattern under oxic conditions between the regular and NH4Cl-free cultures.

**Figure 4:**
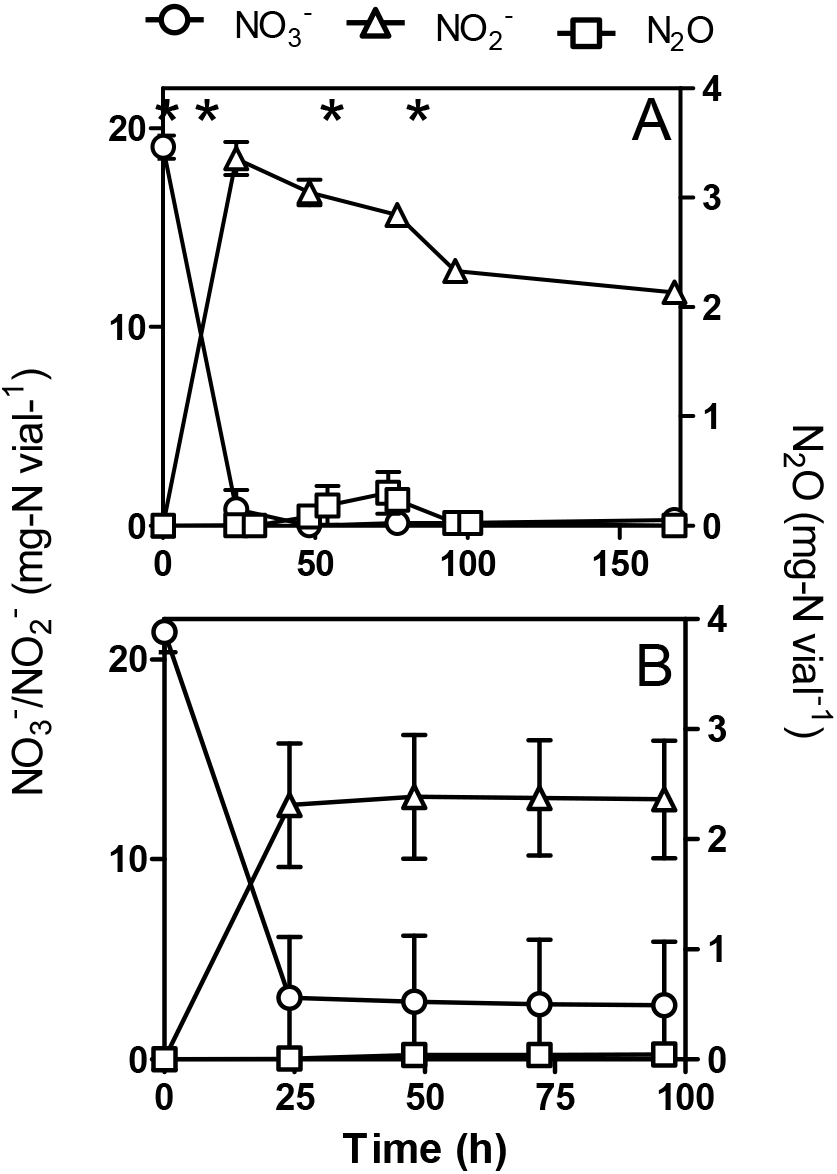
Influence of ammonium on NO3^−^ and NO2^−^ consumption, and N_2_O production and consumption by *Methylophaga nitratireducenticrescens* JAM1. Strain JAM1 was cultured under anoxic (A) or oxic (B) conditions with NO_3_^−^ (22 mg-N vial^-1^) in NH4Cl-free 1403 medium. NO_3_^−^, NO_2_^−^ and N_2_O concentrations were measured over different time intervals. The results are derived from triplicate cultures. In panel A, asterisks denote the sampling times used for RNA extraction (see Figure 6). Data represent mean values ± SD (n=3).

Under anoxic NH4Cl-free conditions, full nitrate reduction (19.1 ± 0.6 mg-N vial^-1^) occurred within 48 h (Fig. 4A). The N_2_O production and consumption profile found was similar to that observed in regular cultures (Fig. 3A), though lower N_2_O concentrations were detected during the accumulating phase. The nitrite level reached 18.5 ± 0.8 mg-N vial^-1^ after 24 h and then slowly decreased to 12.8 ± 0.5 mg-N vial^-1^ after 96 h. Nitrogen assimilation by the biomass and the production of N_2_O and its reduction to N2 could account for the difference in nitrogen mass balance (32.7 ± 2.5%).

Unlike the cultures in regular medium (Fig. 3B), NO_3_^−^ (21.3 ± 1.0 mg-N vial^-1^) was not completely reduced under oxic NH_4_Cl-free conditions, and it stopped after 24 h at 2.9 ± 2.7 mg-N vial^-1^ (Fig. 4A). In conjunction with NO_3_^−^ reduction, NO_2_^−^ levels stopped accumulating at 13.0 ± 2.6 mg-N vial^-1^ after 24 h. N_2_O was observed after 48 h of incubation (Fig. 4A), after which it slowly accumulated and reached a concentration of 0.043 ± 0.048 mg-N vial^-1^. This level is 7 times lower than that of the regular culture medium (Fig. 3B). Nitrogen assimilation by the biomass under oxic conditions could account for the difference in nitrogen mass balance (25.5 ± 4.3%) found between the initial concentration of NO_3_^−^ and residual concentrations of NO_3_^−^, NO_2_^−^ and N_2_O (13.9%, 60.4% and 0.20% of the N input, respectively). In addition, N_2_O production and consumption could have reached an equilibrium and loss of nitrogen would occur by N2 production.

To assess whether N_2_O could have been generated through NH_4_^+^, strain JAM1 was cultured under anoxic conditions with 22 mg-N vial^-1^ NO_3_^−^, 20.7 mg-N vial^-1 15^NH_4_^+^, and acetylene to prevent the reduction of N_2_O to N2. Unfortunately, strain JAM1 cannot be cultured without NO_3_^−^ under anoxic conditions, and growth is inhibited in NO_2_^−^-amended cultures (8). If NH_4_^+^ is involved in N_2_O production, high proportion of labelled N_2_O is expected. If NH_4_^+^ is not involved in N_2_O production, we expected the production of labeled N_2_O to be derived from ^15^NO_3_^−^ naturally present in NaNO_3_^−^ at a natural ^15^N/^14^N isotopic ratio of 0.0036765. In the ^15^NH_4_^+^-amended cultures, the ^45^[N_2_O]/^44^[N_2_O] and ^46^[N_2_O]/^44^[N_2_O] ratios measured were 0.008 and 0.0165, respectively, with an ^15^N/^14^N isotopic ratio of 0.020418. As a control, strain JAM1 cultured under anoxic conditions with ^15^NO_3_^−^ in NH4Cl-free medium with acetylene showed, as was expected, all N_2_O recovered in ^46^[N_2_O]. Because low ^15^N/^14^N isotopic ratio were found in the ^15^NH_4_^+^-amended cultures, our results suggest that N_2_O do not proceed through NH_4_^+^.

### No reduction by M. nitratireducenticrescens JAM1

To verify NO reduction by strain JAM1, N_2_O generation was monitored in cultures without NO_3_^−^ and supplemented with sodium nitroprusside hypochloride (SNP) used as an NO donor (Fig. 5). Because N_2_O is quickly reduced under anoxic conditions but accumulates under oxic conditions, these assays were performed under oxic conditions. N_2_O started to accumulate in both 2 mM and 5 mM SNP-supplemented media after 24 h of incubation, reaching 7.9 ± 0.5 μg-N vial^-1^ and 14.5 ± 0.4 μg-N vial^-1^, respectively, after 168 h. No N_2_O production was observed in strain JAM1 cultures without SNP or in the controls with non-inoculated culture medium supplemented with SNP or inoculated with autoclaved biomass.

**Figure 5:**
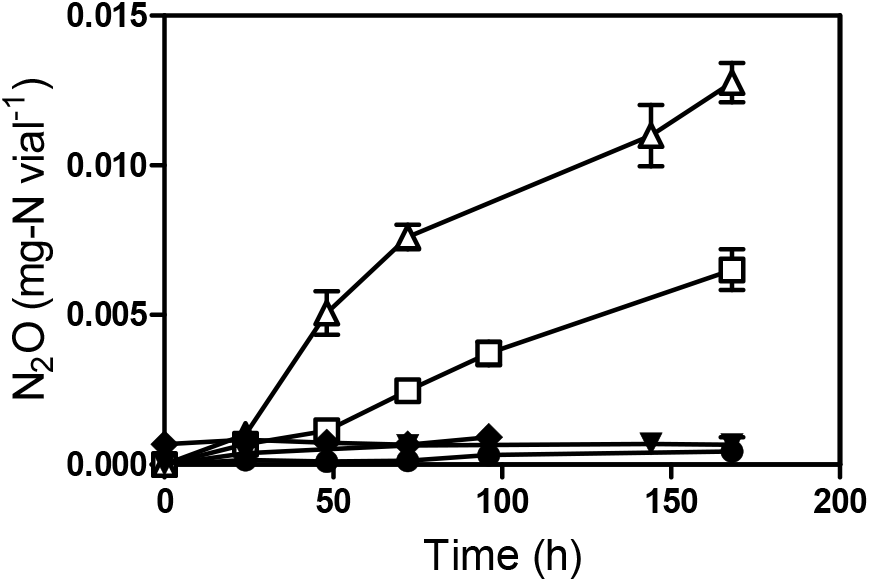
Reduction of NO to N_2_O by *Methylophaga nitratireducenticrescens* JAM1. Strain JAM1 was cultured under oxic conditions without NO_3_^−^ and with 2 mM (square), with 5 mM (triangle), or with no (circle) sodium nitroprusside. N_2_O concentrations were measured over different time intervals. Controls with 5 mM SNP in non-inoculated culture medium (reverse triangle) and in culture medium inoculated with autoclaved biomass (diamond) were also performed. Data represent mean values ± SD (n=3).

### Role of Nar systems in NO/N_2_O production

In the absence of NirK or NirS, N_2_O could have been generated via NO by the Nar system (see discussion). We used the JAM1 Δ*narG1narG2* double mutant, which lacks functional Nar-type nitrate reductases and which cannot grow under anoxic conditions (13). Strain JAM1 and the JAM1 Δ*narG1narG2* were cultured with 16.8 mg-N vial^-1^ NO_3_^−^ under oxic conditions. The growth of strain JAM1 and the mutant was similar (13). After 96 h of incubation, strain JAM1 completely reduced NO_3_^−^ to NO_2_^−^ and produced 0.14 mg-N vial^-1^ of N_2_O (Table 1). As was expected, NO_3_^−^ was not reduced, and NO_2_^−^ was not produced by JAM1 Δ*narG1narG2.* Contrary to the wild type strain, the mutant did not produce N_2_O.

**Table 1:** Production of N_2_O by strain JAM1 and the JAM1Δ*narG1narG2* double mutant.

| Strain | Conditions | Nitrate | Nitrite | N<sub>2</sub>O |
| --- | --- | --- | --- | --- |
| JAM1 | A | 0.17(0.06) | 16.6(0.7) | 0.14(0.01) |
| JAM1ΔnarG1G2 | A | 17.1(0.1) | 0.22(0.22) | 0.004(0.002) |
| JAM1 | B | 0 | 4.25(0.09) | 0.11(0.03) |
| JAM1ΔnarG1G2 | B | 0 | 4.87(0.39) | 0.18(0.02) |

Concentrations of nitrate, nitrite and nitrous oxide (mg-N vial<sup>-1</sup>) were measured after 96 h (OD<sub>600nm</sub> ~ 1.2) of incubation in strain JAM1 and JAM1ΔnarG1narG2 cultured under oxic conditions with (A) 16.8 mg-N vial<sup>-1</sup> NO<sub>3</sub><sup>-</sup> added at T0h or (B) 4.7 mg-N vial<sup>-1</sup> NO<sub>2</sub><sup>-</sup> added at T24h. The results are derived from triplicate cultures. Data represent mean values (SD) (n=3).

The influence of NO_2_^−^ was also tested. As the toxicity of NO_2_^−^ has been attested from 0.36 mM (0.2 mg-N vial^-1^) (8), strain JAM1 and the mutant were cultured without NO_3_^−^ under oxic conditions to allow for biomass growth. After 24 h, 4.7 mg-N vial^-1^ NO_2_^−^ was added to the cultures, which were incubated for another 72 h. Strain JAM1 and the mutant produced 0.11 mg-N vial^-1^ and 0.18 mg-N vial^-1^ of N_2_O, respectively, reflecting N_2_O concentrations produced by strain JAM1 under oxic conditions with NO_3_^−^ (Table 1). Our results show that NO_2_^−^ and not NO_3_^−^ is directly involved in N_2_O production, and the Nar systems are not involved in N_2_O production via NO.

### Indirect detection of NO production by M. nitratireducenticrescens JAM1

We assessed whether variations in the expression levels of denitrification genes correlate with the N_2_O production and consumption cycles of strain JAM1 cultures. Strain JAM1 was cultured in NH4Cl-free medium with 22 mg-N vial^-1^ NO_3_^−^ under anoxic conditions. RNA was extracted from cells harvested over four different phases (Fig. 4): 1) at T0 for the pre-cultures, 2) during the growth phase with nitrate reduction and no N_2_O production, 3) during the N_2_O-production phase, and 4) during the N_2_O-consumption phase. The relative transcript levels of c*norB1*, c*norB2* and *nosZ,* which encode the catalytic subunits of the corresponding NO and N_2_O reductases, and *nnrS*, were measured by RT-qPCR. *nnrS* encodes a NO-sensitive regulator and was used as an indicator of the presence of NO in the cultures. c*norB1* and *nnrS* expression patterns observed were similar, with the highest expression levels observed during the N_2_O-production phase (Fig. 6), which suggests the presence of NO. The c*norB2* expression level remained stable except during the initial phase, when it was significantly lower. With the exception of that of the pre-cultures, the c*norB2* transcript level was always lower than those detected for *cnorB1*. No significant changes in *nosZ* expression levels were observed over the four phases.

**Figure 6:**
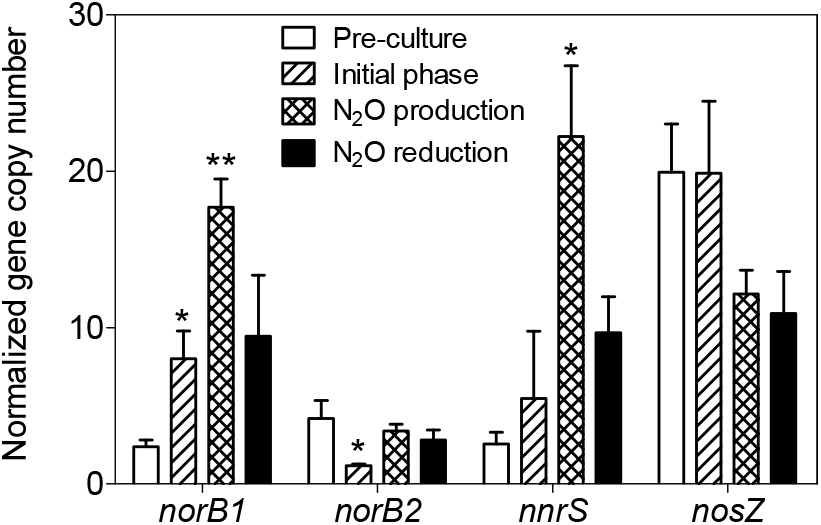
Relative transcript levels of *cnorB1, cnorB2, nnrS* and *nosZ*. Strain JAM1 was cultured under anoxic conditions in NH4Cl-free 1403 medium with 22 mg-N vial^-1^ NO_3_^−^. Growth patterns were similar to those shown in Figure 1B under the same conditions with regular 1403 medium. Samples were drawn from the pre-culture and during the growth phase (no N_2_O production), N_2_O-production phase, and N_2_O-consumption phase (see *Figure 4D*), from which total RNA was extracted. Gene expression levels of *cnorB1, cnorB2, nnrS* and *nosZ* were measured by RT-qPCR and were reported as the gene copy number per copy of *dnaG* (reference gene). Student's *t*-tests were performed for each phase to draw comparisons to the pre-culture phase. *: 0.05<P<0.01; **: 0.01<P<0.001. The results are derived from triplicate cultures from different inoculums. Data represent mean values ± SD (n=3).

## Discussion

Our results show that *M. nitratireducenticrescens* JAM1 can consume NO and N_2_O via the mechanism of reduction of NO to N_2_O and then to N2 as predicted by the genome sequence (Fig. S1) (9, 13). The N_2_O-amended cultures yielded equivalent biomass results to those of the NO_3_^−^-amended cultures as predicted by the respiratory electron transport chains of the denitrification pathway (15). Therefore, in addition of reducing nitrate, strain JAM1 has another respiratory capacity under anoxic conditions by reducing N_2_O for its growth. As observed with nitrate reduction, NO and N_2_O reduction can occur under oxic conditions, reinforcing the lack of a functional oxygen regulation response in strain JAM1 (see Discussion in 13).

N_2_O production was observed in NO_3_^−^- amended cultures either under oxic or anoxic conditions when NO_2_^−^ was accumulating. This production represented up to 6% of N-input in the anoxic cultures, and NO_2_^−^ was shown to be the key element of this production process. Because, we showed that the NO reductase activities were carried out in strain JAM1 cultures, the N_2_O could originate from NO production despite the absence of gene encoding NirS or NirK. Intermediate NO creates problems as this molecule is highly toxic to microorganisms, inducing nitrosative stress in cells (5). Reducing NO is a key step in denitrification and is closely regulated by various sensors and regulators. NnrS is involved in cell defense against nitrosative stress and is positively regulated by the presence of NO (16-18). Therefore, *nnrS* expression reflects NO concentrations in a medium and was used as a marker of NO presence. The higher expressions of *nnrS* found during N_2_O production strongly suggest that NO is produced during this phase. This correlates with higher expressions of *cnorB1,* which can also be regulated by NO-sensitive regulators such as NnrS or NorR (19). Moreover, the increase in *cnorB1* expression found can be directly linked to observed N_2_O production levels. During the N_2_O-consumption phase, only c*norB1* expression decreased and could have changed the balance between N_2_O production and consumption. As *nosZ* expression is mainly reduced by the presence of O2, the stable expression of this gene under constant anoxic conditions was expected (19). Interestingly, c*norB2* expression was not affected by the presence of NO, unlike *cnorB1.* This suggests a putatively different regulation mechanism for this gene like those observed for *narG1* and *narG2* in a previous study (13). Finally, the succession of different phases of N_2_O production/consumption correlates with the presence of NO through an increased expression of *nnrS,* which strongly suggests that NO is an intermediate in N_2_O production in strain JAM1.

Other nitrate respiring bacteria that lack NirK or NirS have been shown to be N_2_O producers (20-22). For instance, *Bacillus vireti* contains three denitrification reductases (Nar, qCuANor, N_2_OR) and lacks, like *M. nitratireducenticrescens* JAM1, gene encoding NirK or NirS (23). This bacterium also produces NO and N_2_O in anaerobic, NO_3_^−^-amended TSB cultures during NO_2_^−^ accumulation. NO was shown to originate from chemical decomposition of NO_2_^−^ (6) and from an unknown biotic reaction. In our study, the abiotic control of the Methylophaga 1403 medium amended with NO_3_^−^ and NO_2_^−^ did not show N_2_O production. Furthermore, no N_2_O was detected in this medium inoculated with autoclaved biomass (Fig. 5). These results rule out NO/N_2_O production by chemical decomposition of NO_2_^−^ in strain JAM1 cultures. The possible biotic source of NO in absence of NirS or NirK has been studied in *Escherichia coli* (see review by Vine and Cole (24). There are supporting evidence that NO is generated in *E. coli* as a side product during nitrite reduction (i) by the cytoplasmic, NADH-dependent nitrite reductase (NirBD), (ii) by the nitrite reductase NrfAB, and (iii) by NarGHI. Vine et al. (25) showed, with mutants defective in these reductases, that NarGHI is the major enzyme responsible of NO production. However, a small production of NO was still occurring in *narG* mutant, suggesting the involvement of another molybdoprotein. In *M. nitratireducenticrescens* JAM1, the double-knockout mutant JAM1 Δ*narG1narG2,* which lacks the two dissimilatory NO_3_^−^ reductases, was still able to produce N_2_O under oxic conditions at the same level of the wild type when NO_2_^−^ was added to the cultures. These results suggest the two Nar systems are not involved in NO production. The genome of strain JAM1 did not reveal gene encoding NrfAB, but contain a gene cluster encoding a cytoplasmic, NADH-dependent nitrite reductase (CP003390.3; Q7A_2620 and Q7A_2621), which may be the source of NO (Fig. S1).

The significance of maintaining an incomplete pathway by *M. nitratireducenticrescens* JAM1 is unclear and may depend upon the original habitat and environment, here the denitrifying biofilm. While *M. nitratireducenticrescens* JAM1 serves as an important actor among the microbial community of the marine biofilm in performing optimal denitrifying activities (10, 26), it was thought to participate uniquely in the reduction of NO_3_^−^ to NO_2_^−^. It was previously proposed that NO_2_^−^ reduction to N2 is carried out by *Hyphomicrobium nitrativorans* NL23, the second most represented bacterium in the biofilm (12, 27). Its capacity to reduce NO and N_2_O and to grow on N_2_O suggests that *M. nitratireducenticrescens* JAM1 may participate in the reduction of NO and N_2_O during denitrification in the biofilm. Although our culture assays were performed with high levels of NO_3_^−^ (37 mM), which is rarely exceeds a value of 0.7 mM in natural environments (28), similar levels can be reached in closed-circuit systems like the seawater aquarium tank located in the Montreal Biodome, where NO_3_^−^ levels reached up to 14 mM (29). Rissanen et al. (30) observed also the combination of *Methylophaga* spp. and *Hyphomicrobium* spp. in the fluidized-bed type denitrification reactors treating the recirculating seawater of the public fish aquarium SEA LIFE at Helsinki, Finland. Although, this study provided no indication of the denitrification pathway in these *Methylophaga* and *Hyphomicrobium,* it reinforces the importance of the natural combination of these two genera in marine denitrification environment.

## Conclusions

*M. nitratireducenticrescens* JAM1 is one of few isolated marine methylotrophic bacterial strains to exhibit anaerobic respiratory capacities by reducing NO_3_^−^ to NO_2_^−^ and, as reported here, by reducing N_2_O to N2. It can also generate N_2_O via NO by an unknown biotic system. Very few marine denitrifying bacteria have been isolated from recirculating marine systems (31-34). No previous studies have generated genetic information related gene arrangement or expression on these bacteria. Based on substantial data accumulated on the genome, gene arrangement and gene expression of denitrification and on methylotrophy, *M. nitratireducenticrescens* JAM1 can serve as a model for studying such activities in marine environments. Finally, our results enable a better understanding of the ecophysiological role of *M. nitratireducenticrescens* JAM1 in the original biofilm developed in the denitrification reactor of a closed-circuit marine aquarium.

## Materials and Methods

### Bacterial growth conditions

*M. nitratireducenticrescens* JAM1 and the JAM1 Δ*narG1narG2* double mutant were cultured in the American Type Culture Collection (ATCC, Manassas, VA, USA) *Methylophaga* medium 1403 (9, 13). When required, NO_3_^−^ (NaNO3) or NO_2_^−^ (NaNO2) (Fisher Scientific Canada, Ottawa, ON, Canada) were added to the medium. Medium (40 or 60 mL) was dispensed into 720-mL bottles (680- or 660-mL head space) that were sealed with caps equipped with septum and which were then autoclaved. After autoclaving, the following filter-sterilized solutions were added to the bottles (40 mL volume): 120 μL methanol (final concentration 0.3% [vol/vol]; 74.3 mM), 800 μL solution T [per 100 mL: 0. 7 g KH2PO4, 10 g NH4Cl, 10 g Bis-Tris, 0.3 g ferric ammonium citrate (pH 8)], 400 μL Wolf’s mineral solution (pH 8) (ATCC), and 40 μL vitamin B12 (stock solution 0.1 mg/mL). The Wolf mineral solution is composed of (per liter) 0.5 g EDTA, 3.0 g MgSO4.7H2O, 0.5 g MnSO4.H2O, 1.0 g NaCl, 0.1 g FeSO4.7H2O, 0.1 g Co(NO3)2.6H2O, 0.1 g CaCl2 (anhydrous), 0.1 g ZnSO4.7H2O, 0.010 g CuSO4.5H2O, 0.010 g AlK(SO4)2 (anhydrous), 0.010 g H3BO3, 0.010 g Na2MoO4.2H2O, 0.001 g Na2SeO3 (anhydrous), 0.010 g Na2WO4.2H2O, and 0.020 g NiCl2.6H2O. The final concentration of ammonium (NH_4_^+^) in the *Methylophaga* 1403 medium was measured as 21 mg-N vial^-1^ (20.9 mg-N vial^-1^ from NH4Cl and 0.1 mg-N vial^-1^ from ferric ammonium citrate). The amount of NO_3_^−^ carried by the Wolf mineral solution (0.0038 mg-N vial^-1^) was deemed negligible. For the anoxic cultures, bottles were flushed with nitrogen gas (N2, purity >99.9%; Praxair, Mississauga, ON, Canada) or argon (purity 99.9%, Praxair) for 20 min prior to autoclaving. When necessary, N_2_O (purity 99.9%, Praxair) and acetylene (10% [vol/vol] of headspace; Praxair) were injected into the headspace before autoclaving. Acetylene is an inhibitor of nitrous oxide reductase and has been extensively used in N_2_O studies to observe N_2_O production in cells (35). Inoculums were made from fresh culture cultivated under oxic conditions without NO_3_^−^ to reach an optical density (OD600) of 0.025. Culture bottles were incubated at 30°C in the dark. For oxic cultures, bottles were shaken at 150 rpm.

The capacity for strain JAM1 to reduce NO was tested with sodium nitroprusside (sodium nitroprusside hypochloride ([SNP]; purity S 99.0%, Sigma-Aldrich, St. Louis, MO, USA) as the NO source. Strain JAM1 was cultured in *Methylophaga* 1403 medium under oxic conditions without NO_3_^−^ for 24 h. The cells were then centrifuged (8,000 g 5 min) and dispersed into fresh medium supplemented with 2 mM, 5 mM, or no SNP. Culture medium with 5 mM SNP and no biomass was also used as a control. Cultures were incubated under oxic conditions, and N_2_O production was monitored. To investigate the potential role of NH_4_^+^ in N_2_O production, NH4Cl-free cultures were employed under oxic and anoxic conditions using solution T containing no NH4Cl. Prior to inoculation, cells from start-up cultures were centrifuged and rinsed three times with saline solution to remove any residual traces of NH_4_^+^.

Bacterial growth was monitored by spectrophotometry (OD600). Bacterial flocs were dispersed with a Potter-Elvehjem homogenizer prior to measurement. Oxygen concentrations in the headspace were monitored in cultures under oxic conditions by gas chromatography using a temperature conductivity detector (7890B series GC Custom, SP1 option 7890-0504/0537; Agilent Technologies, Mississauga, ON, Canada). Although vials were capped in the oxic cultures, O2 concentrations in the headspace (680 ml) did not significantly decrease (T0 h = 20.4 ± 0.3%; T100 h = 19.7 ± 0.9%).

### ^15^N-labeling of N_2_O

Strain JAM1 cultures were made with 22 mg-N vial^-1^ Na^15^NO3 (Sigma-Aldrich) in NH4Cl-free medium or with 22 mg-N vial^-1^ Na^14^NO3 and 20.7 mg-N vial^-1 15^NH4Cl (Sigma-Aldrich). Both cultures were used under anoxic conditions, and 10% (vol/vol) acetylene was added to allow N_2_O to accumulate. Cultures were made in triplicate. After 14 days of incubation, the headspace of each replicate was pooled, and 100 mL of the gaseous phase was sampled in Tedlar bags. N_2_O-isotope measurements were performed at the Environmental Isotope Laboratory (Earth & Environmental Sciences; University of Waterloo, ON, Canada) via Trace Gas-GVI IsoPrime-Isotope Ratio Mass Spectrometry (TG-IRMS). ^45^[N_2_O]/^44^[N_2_O] and ^46^[N_2_O]/^44^[N_2_O] ratios were calculated according to the peak intensity measured for ^46^[N_2_O], ^45^[N_2_O] and ^44^[N_2_O]. The ^15^N/^14^N isotopic ratio was derived from the previous results from Eq. 3.

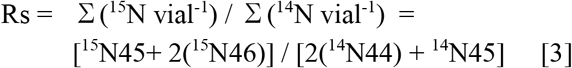

where Rs is the sample isotopic ratio. Calculated from the ^45^[N_2_O]/^44^[N_2_O] and ^46^[N_2_O]/^44^[N_2_O] isotopic ratios, ^14^N45 is the quantity of ^14^N in ^45^[N_2_O], ^15^N45 is the quantity of ^15^N in ^45^[N_2_O], ^14^N44 is the quantity of ^14^N in ^44^[N_2_O] and ^15^N46 is the quantity of ^15^N in ^46^[N_2_O]. We considered the isotope fractionation by denitrification enzymes as negligible in our calculations (delta values ranging from -10 %o to -40 %o) (36).

### Measurements of nitrogenous compounds

NO_3_^−^ and NO_2_^−^ concentrations were determined by ion chromatography using the 850 Professional IC (Metrohm, Herisau, Switzerland) with a Metrosep A Supp 5 analytical column (250 mm × 4.0 mm).

N_2_O and N2 concentrations were determined by gas chromatography. Headspace samples (10 mL) were collected using a Pressure Lok gastight glass syringe (VICI Precision Sampling Inc., Baton Rouge, LA, USA) and were injected through the injection port of a gas chromatograph equipped with a thermal conductivity detector and electron-capture detector (7890B series GC Custom, SP1 option 7890-0504/0537; Agilent Technologies). The reproducibility of the N_2_O was assessed before each set of measurements was conducted via the repeated analysis of certified N_2_O standard gas with standard deviations <5%. N_2_O standards (500 ppmv and 250 ppmv) were created based on dilutions from the 10,000 ppmv N_2_O stock standard. The 10,000 ppmv stock standard was obtained by injecting 1% pure N_2_O (Praxair) into a 720 mL gastight bottle. The detection limit of the N_2_O was set to <10 ppbv, corresponding to the 0.3 nmol/vial composition of our bioassays. No significant N_2_O production patterns were observed through our blank experiments involving sterile media and empty glass bottles. The total quantity of N_2_O in the culture bottle (aqueous phase and headspace) (X_N2O_ in μmole vial^-1^) was calculated according to Eq. 4.

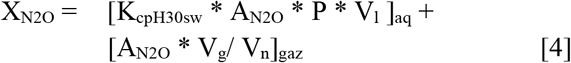

where AN_2_O: the N_2_O mixing ratio measured in the headspace (ppmv/10^6^; no unit); P: 1 atm; V1 and Vg: volume of the aqueous (0.04 or 0.06 L vial^-1^) and gaseous phases (0.68 or 0.66 L vial^-1^), respectively; and Vn: molar volume [RT (gas constant): 0.08206 L atm K^-1^ mol^-1^ * 303K = 24.864 L mol^-1^]. K_H30sw_ is the corrected Henry's constant for seawater at 30°C (0.01809 mol L^-1^ atm^-1^) according to Weiss and Price (1980). XN_2_O was then converted (eq.5) in an mg-N vial^-1^ for an easier calculation of mass balances using the other nitrogenous compounds:

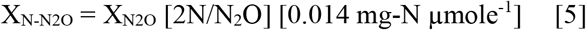

The reproducibility of the N2 was assessed before each set of measurements was made by a repeated analysis of N2 (purity >99.99%, Praxair) diluted in a 720 mL gastight bottle (0 and 500 ppmv) flushed with argon (purity >99.99%, Praxair). The total quantity of N2 in the culture bottles was only considered for the headspace, as the quantity of dissolved N2 in the aqueous phase was considered to be negligible in our experimental design based on Henry’s constant (0.0005 mol L^-1^ atm^-1^) and was thus calculated according to Eq. 6.

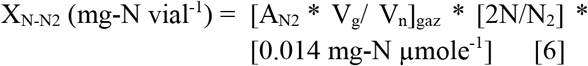

### RNA extraction

Anoxic cultures of strain JAM1 were created in an NH4Cl-free 1403 medium supplemented with 22 mg-N vial^-1^ NO_3_^−^. Cells were harvested at specific times, and RNA was immediately extracted using the PureLink RNA mini kit (Ambion Thermo Fisher Scientific, Burlington, ON, Canada). RNA extracts were treated twice with TurboDNase (Ambion), and RNA quality was verified by agarose gel electrophoresis. The absence of remaining DNA was checked via the end-point polymerase chain reaction (PCR) amplification of the 16S rRNA gene using RNA extracts as the template.

### Gene expression

cDNAs samples were generated from the RNA using hexameric primers and the Reverse Transcription System developed by Promega (Madison, WI, USA) with 1 μg of RNA. Real-time quantitative PCR (qPCR) assays were performed using the Faststart SYBR Green Master (Roche Diagnostics, Laval, QC, Canada) according to the manufacturer’s instructions. All reactions were performed in a Rotor-Gene 6000 real-time PCR thermocycler (Qiagen Inc. Toronto, ON, Canada), and each reaction contained 25 ng of cDNA and 300 nM of primers (Table 2). Genes tested included *cnorB1, cnorB2, nosZ* and *nnrS.* PCR began with an initial denaturation step of 10 min at 95°C followed by 40 cycles of 10 s at 95°C, 15 s at 60°C, and 20 s at 72°C. To confirm the purity of the amplified products, a melting curve was performed by increasing the temperature from 65°C to 95°C at increments of 1°C per step with a pause of 5 s included between each step. All genes for each sample and standard were tested in a single run. The amplification efficiency level was tested for each set of primer pairs by qPCR using a dilution of strain JAM1 genomic DNA as the template. The amplification efficiencies for all primer pairs varied between 0.9 and 1.1. The copy number of each gene was calculated according to standard curves using dilutions of strain JAM1 genomic DNA. To normalize the gene expression of the different growth phases, results were expressed as copy numbers per *dnaG* copy numbers for each sample. In accordance with previous studies (13), *dnaG* generated the least variability of the reference genes tested (*dnaG, rpoD* and *rpoB)* in strain JAM1. *dnaG* encodes for a DNA primase and is present in one copy in strain JAM1 genomes. RNA extraction and qPCR were performed with three independent biological replicates. The significance of differential expression levels was tested for each phase against the pre-culture phase via Student’s t-test.

**Table 2:**
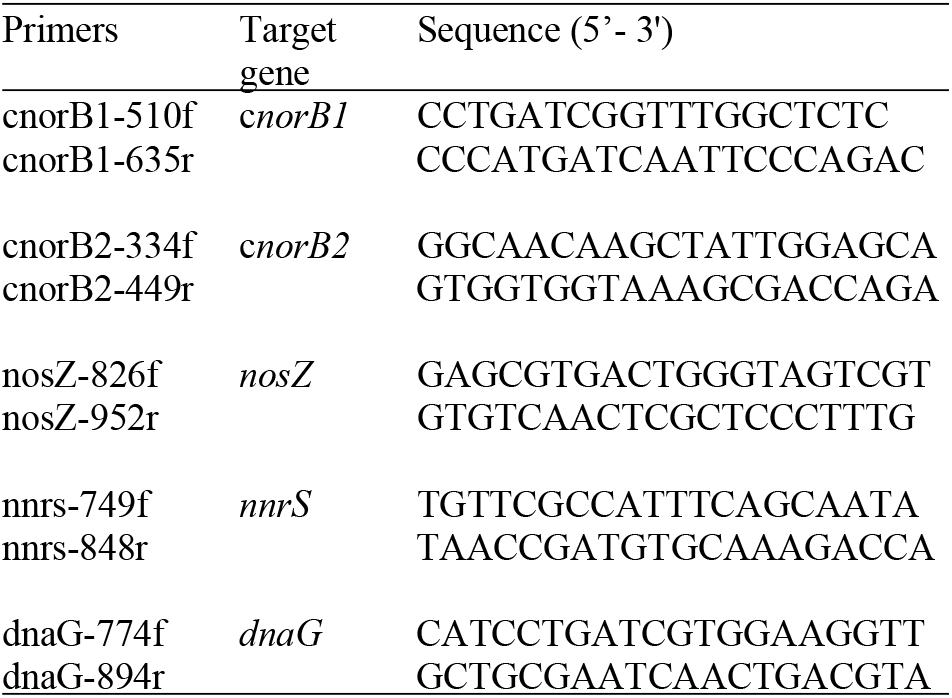
Primers used for RT-qPCR.

## Acknowledgments

We thank Karla Vasquez for her technical assistance. This project was financially supported by a grant to RV from the Natural Sciences and Engineering Research Council of Canada (#155558).

---

**Figure S1:**
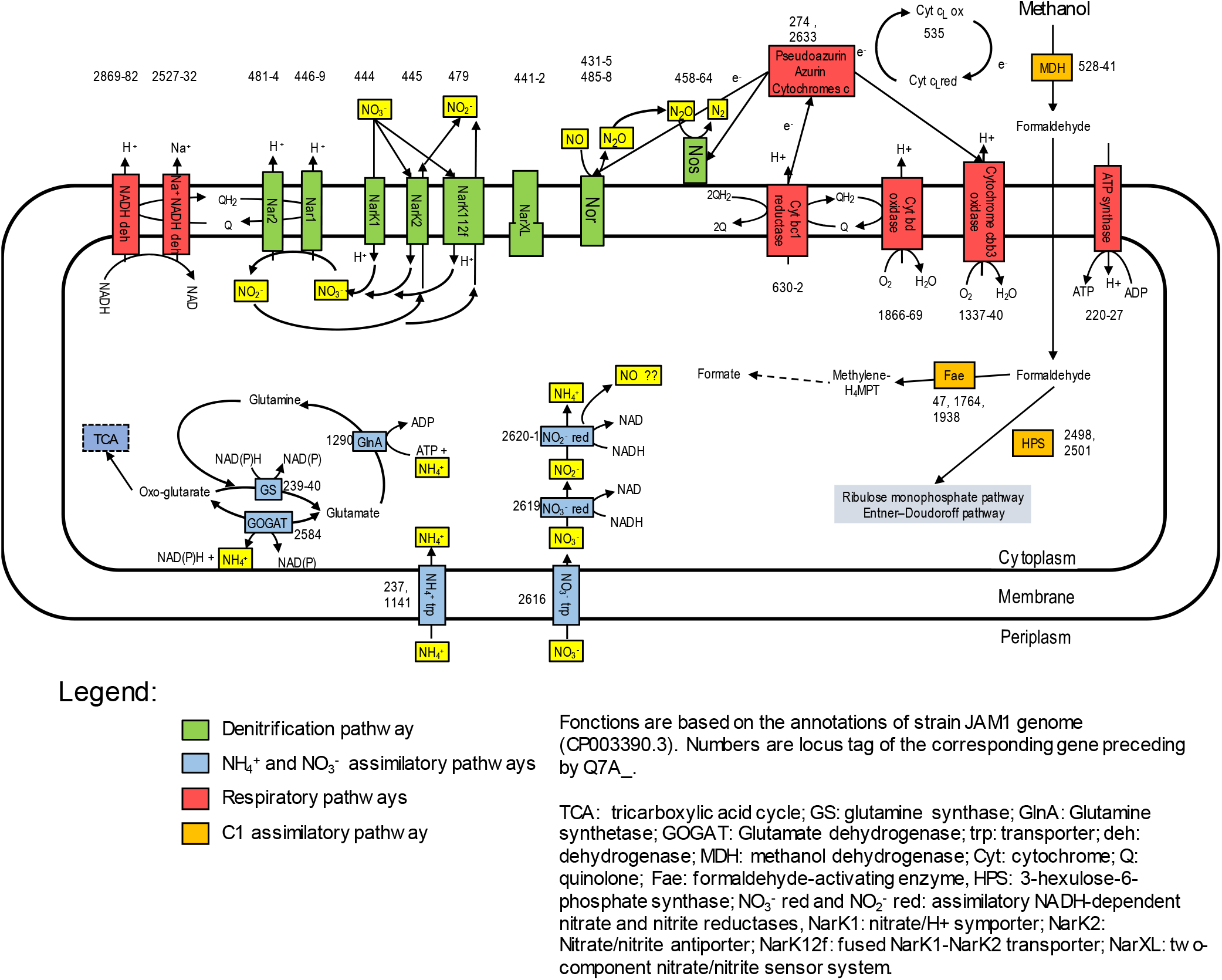
Schematic of the denitrification pathway and the ammonium pathways in *Methylophaga nitratireducenticrescens* JAM1.

